# HMP16SData: Efficient Access to the Human Microbiome Project through Bioconductor

**DOI:** 10.1101/299115

**Authors:** Lucas Schiffer, Rimsha Azhar, Lori Shepherd, Marcel Ramos, Ludwig Geistlinger, Curtis Huttenhower, Jennifer B Dowd, Nicola Segata, Levi Waldron

## Abstract

Phase 1 of the NIH Human Microbiome Project (HMP) investigated 18 body subsites of 239 healthy American adults, to produce the first comprehensive reference for the composition and variation of the “healthy” human microbiome. Publicly-available data sets from amplicon sequencing of two 16S rRNA variable regions, with extensive controlled-access participant data, provide a reference for ongoing microbiome studies. However, utilization of these data sets can be hindered by the complex bioinformatic steps required to access, import, decrypt, and merge the various components in formats suitable for ecological and statistical analysis. The *HMP16SData* package provides count data for both 16S variable regions, integrated with phylogeny, taxonomy, public participant data, and controlled participant data for authorized researchers, using standard integrative Bioconductor data objects. By removing bioinformatic hurdles of data access and management, *HMP16SData* enables epidemiologists with only basic R skills to quickly analyze HMP data.

## Introduction

The NIH Human Microbiome Project (HMP) was one of the first large-scale population studies of microbiome variation outside of disease, including healthy American adults aged 18 to 40 and producing a comprehensive reference for the composition and variation of the “healthy” human microbiome (1,2). Raw and processed 16S rRNA (16S) and metagenomic shotgun (MGX) sequencing data can be freely downloaded from the HMP data analysis and coordinating center (DACC) (https://www.hmpdacc.org/hmp/) or analyzed online using the HMP data portal (https://portal.hmpdacc.org/).

However, accessing and analyzing the data with statistical software still involves substantial bioinformatic and data management challenges. These include data import and merging of microbiome profiles with public and controlled-access participant data, integration with phylogenetic trees, potentially mapping microbial and participant identifiers for comparison between 16S and MGX data sets, and accessing controlled participant data.

We thus developed the *HMP16SData* R/Bioconductor package to simplify access to and analysis of HMP 16S data. The design of the package follows our *curatedMetagenomicData* R/Bioconductor package (3), enabling comparative analysis with MGX samples from the HMP and dozens of other studies. *HMP16SData* leverages Bioconductor’s *ExperimentHub* and the *SummarizedExperiment (4)* data class to distribute merged taxonomic and public-access participant data. It provides 16S gene sequencing data for variable regions 1–3 (V13) and 3–5 (V35), with merged participant data, and a function for automated merging of controlled-access participant data to researchers with a project approved by dbGaP. Methods for efficient subsetting and coercion to the *phyloseq* class, which is used by the *phyloseq* package for comparative ecological and differential abundance analyses (5), are also provided. Finally, *HMP16SData* greatly simplifies access to and merging of restricted participant data. These simplifications enable epidemiologists with only basic R skills and limited knowledge of HMP DACC or dbGaP procedures to quickly make use of the HMP data.

## Methods

*HMP16SData* provides data from the HMP 16S compendium as processed through the HMP DACC QIIME pipeline (https://www.hmpdacc.org/HMQCP/) (6). All publicly-available participant data, as obtained from the HMP DACC, is also included and a function provides simplified access to and merging of controlled data from dbGaP for registered researchers. Use of *HMP16SData* begins with one of two functions: V13 (to download data for 16S V13) or V35 (to download data for 16S V35), each of which returns a *SummarizedExperiment* object. Each object contains data/metadata (with specific accessors) as follows: experiment-level metadata (metadata), sample-level metadata (colData), sequencing count data (assay), phylogenetic classification data (rowData), and a phylogenetic tree (metadata). Selection of samples by body site, visit number, and taxonomic hierarchy is straightforward through standard *SummarizedExperiment* or *phyloseq* subsetting methods. Researchers with an approved dbGaP project can optionally use a second function, attach_dbGaP, to attach controlled participant data, prior to coercion to a *phyloseq* object for ecological analyses such as alpha and beta diversity. **Example 1** demonstrates the selection of only stool samples, attachment of controlled participant data, and coercion to a *phyloseq* object. Additional examples are provided in the package vignette and complete function documentation is available in the reference manual.

**Example 1.**
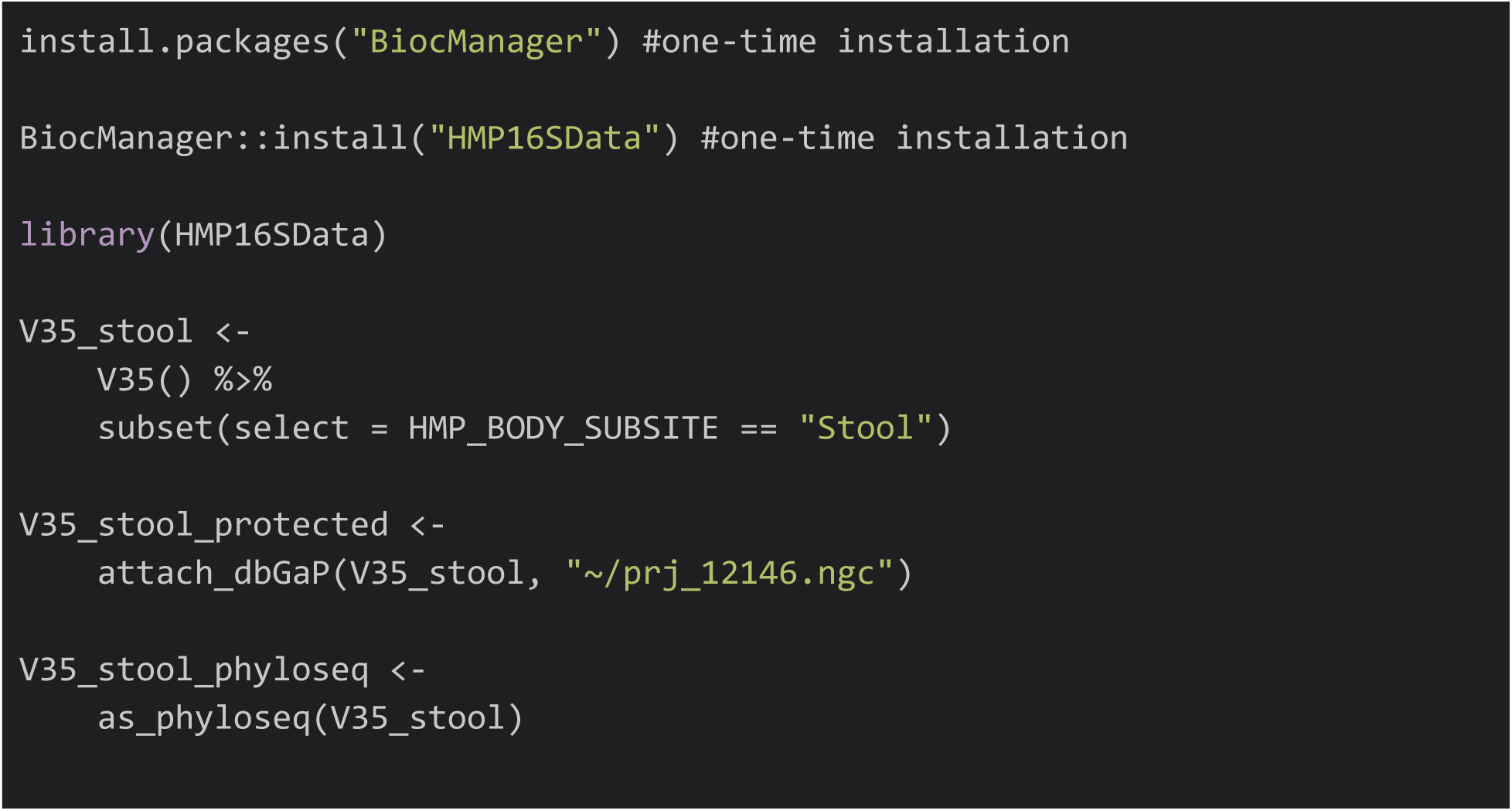
Data access using HMP16SData. This example demonstrates subsetting by body subsite, attaching of controlled participant data from dbGaP, and coercion to a phyloseq class object.

### Controlled-Access Participant Data Analysis

Access to most participant data is controlled, requiring authorization through the National Center for Biotechnology Information (NCBI) Database of Genotypes and Phenotypes (dbGaP), and requires the use of specialized software for download and decryption. Specifically, non-restricted participant data include only visit number, sex, run center, body site, and body subsite – an additional 248 participant data variables are available through dbGaP after project registration. The process involves making a controlled-access application through https://dbgap.ncbi.nlm.nih.gov for HMP Core Microbiome Sampling Protocol A (HMP-A) (phs000228.v4.p1) – see the package vignette for specific details. After project approval, dbGaP provides researchers with a “repository key” that identifies and decrypts controlled-access participant data. The attach_dbGaP function takes the public *SummarizedExperiment* data set provided by *HMP16SData* and the path to the dbGaP repository key as arguments; it performs download, decryption, and merging of controlled participant data, and returns another *SummarizedExperiment* with controlled-access participant data added to its *colData* slot. Internally, attach_dbGaP uses system calls to the NCBI SRA (Sequence Read Archive) Toolkit for download and decryption, and R functionality to load and merge the controlled data. A data dictionary describing the controlled-access participant data variables is incorporated into the package and is accessible by entering data(dictionary).

### Phyloseq Class Coercion

The *phyloseq* package is a commonly used tool for ecological analysis of microbiome data in R/Bioconductor. *HMP16SData* provides a function, as_phyloseq, to coerce its default *SummarizedExperiment* objects to *phyloseq* objects. The resulting objects contain taxonomic abundance count data, participant data, complete taxonomy, and phylogenetic trees, enabling computation of UniFrac (7) and other ecological distances.

## Results

*HMP16SData* provides a total of 7,641 taxonomic profiles from 16S variable regions 1–3 and 3–5 for 239 participants in the HMP, for 18 body subsites and up to three visits. The two variable regions, 1–3 (V13) and 3–5 (V35), identified 43,140 and 45,383 taxonomic clades, respectively, with resolution down to the genus level at a median sequencing depth of 4,428 reads per specimen. These profiles are provided as two Bioconductor *SummarizedExperiment* class objects: V13 and V35 (**Table 1**), which integrate OTU count data, taxonomy, a phylogenetic tree, and public-use participant information. Each object includes both 16S and MGX sample identifiers, enabling mapping and comparison to MGX profiles distributed by our *curatedMetagenomicData* R/Bioconductor package (3). Such a comparison is illustrated in the phylum-level relative abundance plots of matched 16S and MGX sequencing samples in **Figure 1**. Code to reproduce **Table 1** and **Figure 1** are provided in the package vignette documentation along with additional analysis examples.

**Table 1.**
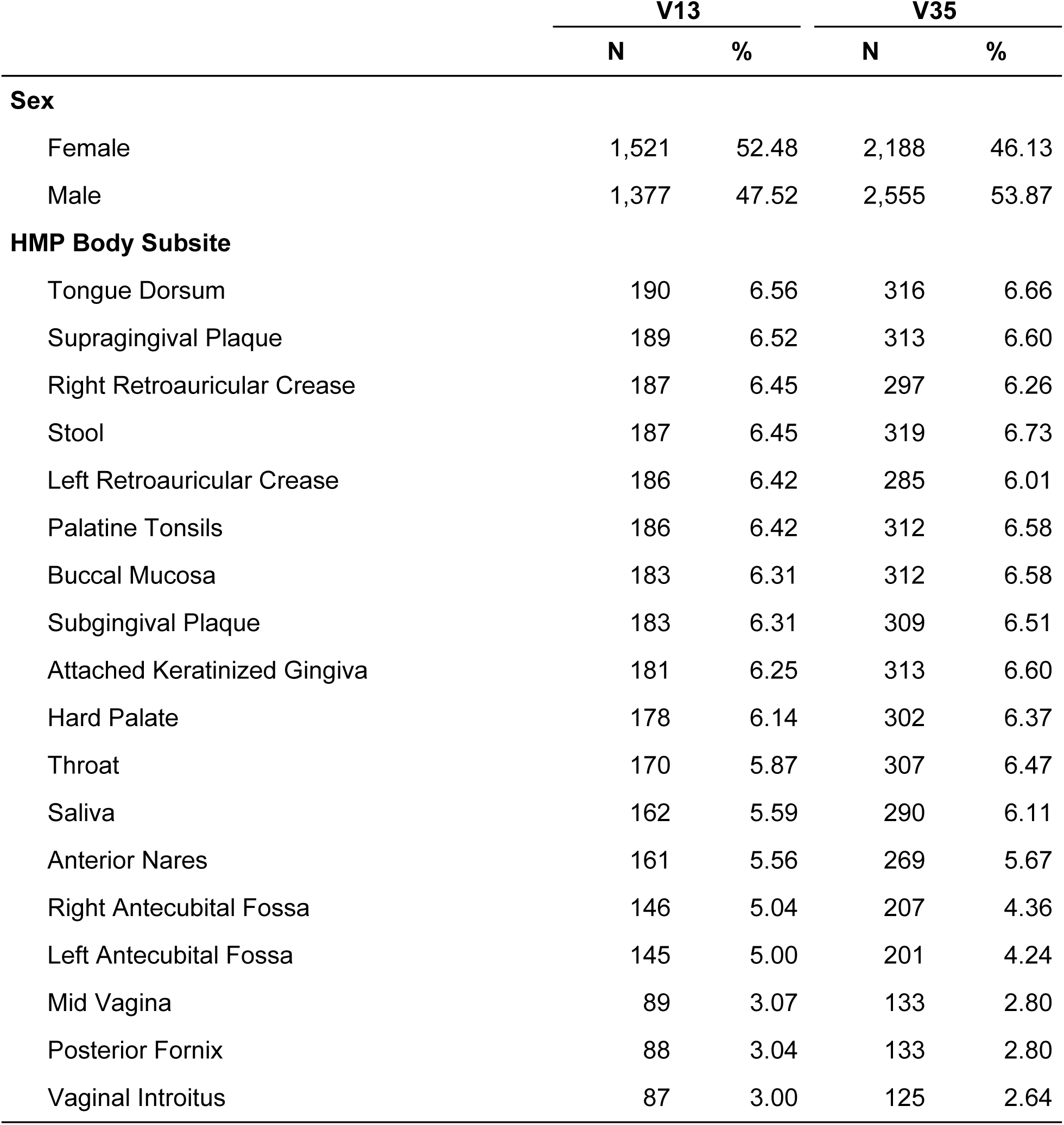
Select characteristics of 16S rRNA (16S) samples for variable regions 1–3 (V13) and 3–5 (V35) available through HMP16SData. All numbers represent samples rather than subjects, given that there are multiple samples per subject.

|  | V13 |  | V35 |  |
| --- | --- | --- | --- | --- |
|  | N | % | N | % |
| **Sex** |  |  |  |  |
| Female | 1,521 | 52.48 | 2,188 | 46.13 |
| Male | 1,377 | 47.52 | 2,555 | 53.87 |
| **HMP Body Subsite** |  |  |  |  |
| Tongue Dorsum | 190 | 6.56 | 316 | 6.66 |
| Supragingival Plaque | 189 | 6.52 | 313 | 6.60 |
| Right Retroauricular Crease | 187 | 6.45 | 297 | 6.26 |
| Stool | 187 | 6.45 | 319 | 6.73 |
| Left Retroauricular Crease | 186 | 6.42 | 285 | 6.01 |
| Palatine Tonsils | 186 | 6.42 | 312 | 6.58 |
| Buccal Mucosa | 183 | 6.31 | 312 | 6.58 |
| Subgingival Plaque | 183 | 6.31 | 309 | 6.51 |
| Attached Keratinized Gingiva | 181 | 6.25 | 313 | 6.60 |
| Hard Palate | 178 | 6.14 | 302 | 6.37 |
| Throat | 170 | 5.87 | 307 | 6.47 |
| Saliva | 162 | 5.59 | 290 | 6.11 |
| Anterior Nares | 161 | 5.56 | 269 | 5.67 |
| Right Antecubital Fossa | 146 | 5.04 | 207 | 4.36 |
| Left Antecubital Fossa | 145 | 5.00 | 201 | 4.24 |
| Mid Vagina | 89 | 3.07 | 133 | 2.80 |
| Posterior Fornix | 88 | 3.04 | 133 | 2.80 |
| Vaginal Introitus | 87 | 3.00 | 125 | 2.64 |

**Figure 1.**
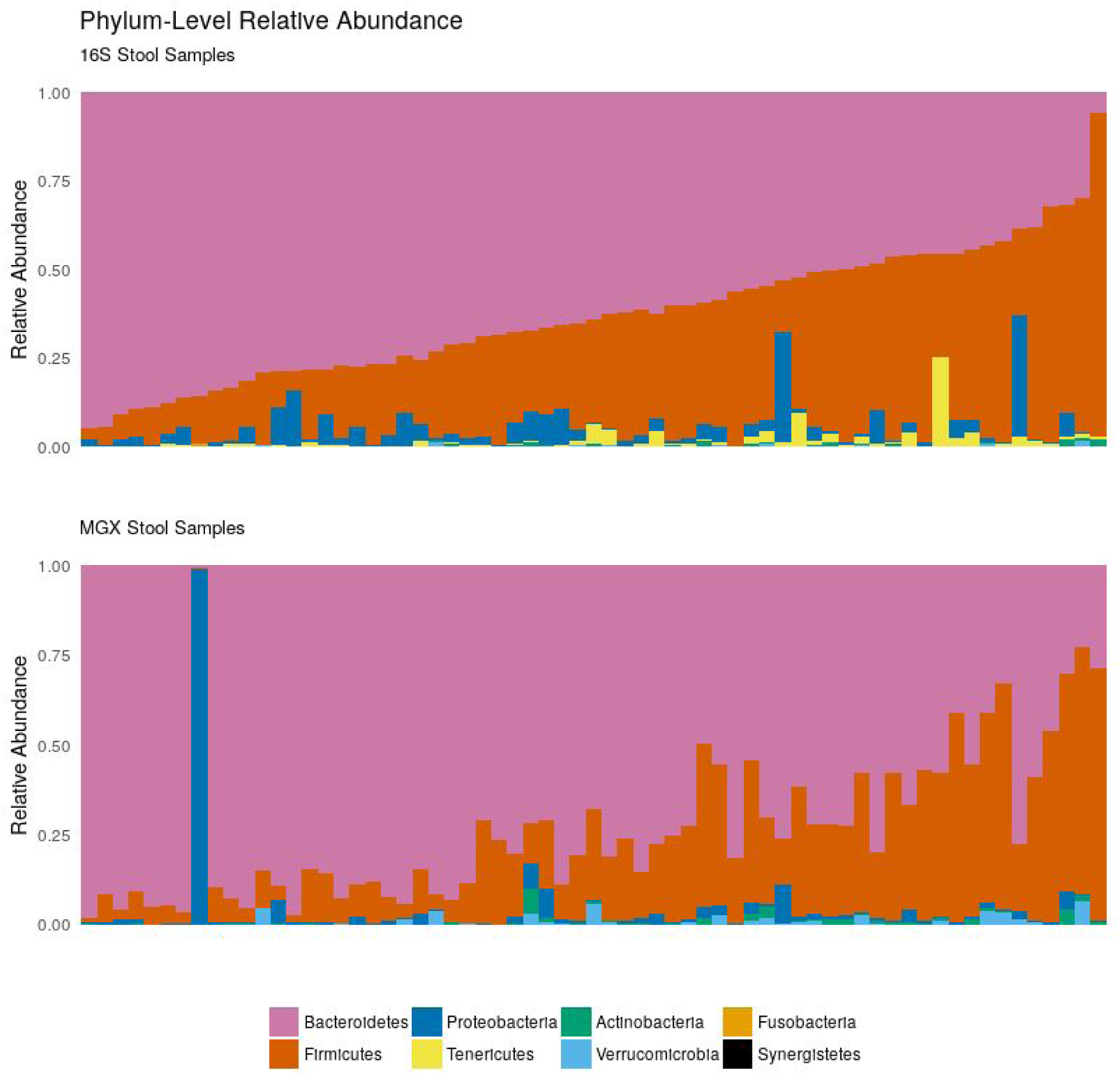
Phylum-level relative abundance of the eight most abundant phyla in matched 16S and MGX samples from the HMP. Samples are ordered in both plots by abundance of Bacteroidetes in 16S samples. Notably, the figure illustrates the Bacteroidetes/Firmicutes gradient with reasonable agreement between the 16S and MGX samples.

## Discussion

The HMP provides a comprehensive reference for the composition, diversity, and variation of the human microbiome in the absence of overt disease, making it a potential control or comparison cohort for many microbiome studies. The R/Bioconductor environment provides an extensive range of operations for data analysis, with documented workflows (8) available for typical microbiome investigations. The *HMP16SData* package thus integrates HMP 16S taxonomic abundance profiles, controlled-access and public participant data, and phylogenetic distances with R/Bioconductor. This greatly reduces the time and bioinformatics expertise required to analyze these data, particularly in the context of additional integrated microbiome population studies. Further, users of other analysis environments can easily export the data to other formats (SAS, SPSS, STATA, etc.) using the haven R package – see the package vignette for specific details (9). We hope this facilitates broader utilization of the data generated by the HMP among epidemiologists, statisticians, and computational biologists.

Some precautions should be noted when using *HMP16SData* in comparative metagenomic analyses. First, studies of the human microbiome are susceptible to batch effects which should be accounted for in making cross-study comparisons, along with other forms of technical variation (10,11). Second, the V13 and V35 data sets are obtained from sequencing different variable regions of the 16S rRNA gene, and provide correlated but different estimates of taxonomic relative abundance (12,13). In the case of the HMP, the samples sequenced in V13 are a subset of those that were sequenced in V35. The two variable regions differ in their ability to distinguish specific microbial clades, e.g. V13 is preferred for discriminating *Streptococci*. Users should choose the better region for their purposes based on the comparison to metagenomic shotgun sequencing provided by Supplemental Figure 3 of the original Human Microbiome Project paper (1) or by the more thorough technical evaluation of the HMP 16S data sets provided by the Jumpstart Consortium (14).

The number of sequence reads per specimen varies by body site and by specimen, from a maximum of 151 thousand reads from a single specimen to a minimum of one read. *HMP16SData* retains all available data, but most analyses should include a quality control step of removing specimens with very low sequencing depth caused by amplification failure or lack of microbial DNA in the specimen. For example, 14% of skin specimens have fewer than one hundred V35 reads, compared to only 4% for V13 reads.

Finally, users of these data should be aware that the HMP data were generated using a legacy platform and processing pipelines. The samples from 15 (male) or 18 (female) body sites of phenotyped adults between the ages of 18 and 40 years who passed a screening for systemic health based on oral, cutaneous and body mass exclusion criteria were sequenced on the Roche 454 FLX Titanium platform (2). The platform is no longer used and the software pipelines, QIIME version 1.3 and mothur version 1.1.8, used to produce OTU tables have since been updated significantly. As such, cross-study analysis of differential abundance or clustering requires care and attention to the biases arising from these differences, particularly when comparing to data produced by more recent sequencing platforms and software pipelines. These data, however, remain a key reference for the general structure of the “healthy” human microbiome.

With these precautions in mind, this resource is intended to provide a key reference data set for the general structure of the “healthy” human microbiome. It enables efficient access to and analysis of the HMP by greatly reducing previous hurdles of data access and management. Finally, while we have simplified the use of the legacy HMP data, we also understand the need to reprocess the raw sequencing data through contemporary bioinformatics tools and will seek to do so in the future.

## Acknowledgements

This research was supported by the National Institute of Allergy and Infectious Diseases (1R21AI121784-01 to J.B.D. and L.W.), the National Cancer Institute (5U24CA180996 to Martin Morgan), the National Institute of Dental and Craniofacial Research (U54DE023798 to C.H.), the National Human Genome Research Institute (R01 HG005220 to Rafael Irizarry), the National Science Foundation (MCB-1453942 and DBI-1053486 to C.H.), and in part, under National Science Foundation Grants CNS-0958379, CNS-0855217, ACI-1126113 to the City University of New York High Performance Computing Center at the College of Staten Island.

The color palette used in **Figure 1** is optimized for color-blind individuals as proposed by Wong (15).

We would like to thank our colleagues Paul J. McMurdie and Susan Holmes, the authors of the *phyloseq* Bioconductor package, whose work has greatly enabled our own (5).

A special thank you to Rodney Middleton for his daily encouragement and desk visits that made this project successful.

## References

1. Human Microbiome Project Consortium. Structure, function and diversity of the healthy human microbiome. Nature. 2012;486(7402):207–214.

2. Human Microbiome Project Consortium. A framework for human microbiome research. Nature. 2012;486(7402):215–221.

3. Pasolli E, Schiffer L, Manghi P, et al. Accessible, curated metagenomic data through ExperimentHub. Nat. Methods. 2017;14(11):1023–1024.

4. Huber W, Carey VJ, Gentleman R, et al. Orchestrating high-throughput genomic analysis with Bioconductor. Nat. Methods. 2015;12(2):115–121.

5. McMurdie PJ, Holmes S. phyloseq: an R package for reproducible interactive analysis and graphics of microbiome census data. PLoS One. 2013;8(4):e61217.

6. Caporaso JG, Kuczynski J, Stombaugh J, et al. QIIME allows analysis of high-throughput community sequencing data. Nat. Methods. 2010;7(5):335–336.

7. Lozupone C, Lladser ME, Knights D, et al. UniFrac: an effective distance metric for microbial community comparison. ISME J. 2011;5(2):169–172.

8. Callahan BJ, Sankaran K, Fukuyama JA, et al. Bioconductor Workflow for Microbiome Data Analysis: from raw reads to community analyses. F1000Res. 2016;5:1492.

9. Wickham H, Miller E. haven: Import and Export “SPSS”, “Stata” and “SAS” Files. 2018.

10. Huttenhower C, Knight R, Brown CT, et al. Advancing the microbiome research community. Cell. 2014;159(2):227–230.

11. Gibbons S, Duvallet C, Alm EJ. Correcting for batch effects in case-control microbiome studies. bioRxiv. 2018;165910.

12. Chakravorty S, Helb D, Burday M, et al. A detailed analysis of 16S ribosomal RNA gene segments for the diagnosis of pathogenic bacteria. J. Microbiol. Methods. 2007;69(2):330–339.

13. Yang B, Wang Y, Qian P-Y. Sensitivity and correlation of hypervariable regions in 16S rRNA genes in phylogenetic analysis. BMC Bioinformatics. 2016;17:135.

14. Jumpstart Consortium Human Microbiome Project Data Generation Working Group. Evaluation of 16S rDNA-based community profiling for human microbiome research. PLoS One. 2012;7(6):e39315.

15. Wong B. Points of view: Color blindness. Nat. Methods. 2011;8:441.

